# Investigating intestinal epithelium metabolic dysfunction in Celiac Disease using personalized genome-scale models

**DOI:** 10.1101/2024.04.29.591234

**Authors:** Chloe V. McCreery, Drew Alessi, Katarina Mollo, Alessio Fasano, Ali R. Zomorrodi

## Abstract

Celiac Disease (CeD) is an autoimmune condition characterized by an aberrant immune response triggered by the ingestion of gluten, which damages epithelial cells lining the small intestine. Small intestinal epithelial cells (sIECs) play a key role in various metabolic processes, including the enzymatic digestion and absorption of nutrients. Although nutritional malabsorption is widely recognized in CeD, the underlying disrupted metabolic processes remain largely undefined. To address this knowledge gap, we constructed personalized gender-specific genome-scale models of sIEC metabolism using transcriptional data from 42 subjects with active CeD, remission CeD, and healthy controls. We computationally simulated these models under a relevant diet for each group of subjects to assess the activity of 59 metabolic tasks essential for sIEC function and to profile metabolite secretion into the bloodstream and intestinal lumen. These investigations revealed significant variations in the activity of 25 metabolic tasks in active and remission CeD models. These tasks impact critical processes integral to sIEC function such as amino acid metabolism, nucleotide synthesis and DNA repair, ATP generation, and oxidative stress regulation. Additionally, we identified 54 metabolites with altered secretion profiles in CeD, encompassing amino acids, vitamins, antioxidants, and fatty acids. Furthermore, we pinpointed 22 FDA-approved drugs that target the genes associated with differentially active metabolic functions whose altered activities adversely affect sIECs in CeD, potentially helping to restore their normal activity. Our study unveils new insights into the metabolic reprogramming of sIECs in CeD, paving the way for therapeutic interventions targeting dysregulated metabolic processes.

## Introduction

The intestinal epithelium is a monolayer of columnar epithelial cells that plays a pivotal role in maintaining the integrity and functionality of the gastrointestinal tract. These cells serve as a critical barrier between the luminal contents of the intestine and the underlying tissues, selectively preventing the permeation of luminal endotoxins, pathogenic microorganisms, and other antigens while allowing the absorption of nutrients, useful microbial products, electrolytes, and water. Additionally, intestinal epithelial cells contribute to mucosal immune regulation and immune tolerance to dietary compounds. However, in the case of inflammatory diseases, the function of intestinal epithelial cells can be profoundly compromised. This can be manifested as a breached gut barrier function ("leaky gut”) allowing the dysregulated passage of endotoxins, pathogens, antigens, and other pro-inflammatory substances into the human body. This in turn can lead to an overactive immune response and subsequently inflammation and disease. One example of such inflammatory conditions is Celiac Disease (CeD), which is the focus of this study.

CeD is an autoimmune disease triggered by the consumption of gluten, a composite protein found in wheat, rye, barley, and other cereal grains. Currently, this disease is estimated to affect over three million Americans and 1.4% of the global population, with a slightly higher prevalence in women (1, 2). CeD has a strong genetic basis with individuals carrying certain human leukocyte antigen (HLA) alleles, namely HLA-DQ2 and HLA-DQ8, being at significantly elevated risk of developing the disease. Chronic exposure to gluten in individuals with genetic predisposition for CeD elicits an abnormal immune response to undigested gluten peptides, particularly those from gliadin, a component of gluten which is resistant to proteolytic digestion.

CeD diagnosis is typically through an intestinal biopsy. However, blood tests measuring tissue transglutaminase (tTG; also known as transglutaminase 2) levels are also used as an effective screening tool in the diagnosis process (3). The autoantigen tTG deamidates undigested gluten peptides. This modification increases the binding affinity of these gluten peptides on antigen-presenting cells expressing on their surface the HLA-DQ2 and/or HLA-DQ8 alleles. These modified gliadin peptides are then presented to T cells, which triggers a strong immune response causing inflammation, damage to the villi and lining of small intestine (i.e., intestinal epithelium), and poor intestinal barrier integrity. This ultimately leads to the establishment of a chronic autoimmune inflammatory process, as well as a range of gastrointestinal symptoms such as abdominal pain, diarrhea, and bloating. Although CeD and its symptoms can be managed by following a strict gluten-free diet, no treatments or drugs exist yet to complement this diet or, even more ambitiously, to return to an unrestricted diet (2, 4).

In addition to a breached gut barrier function, there are other reasons why small intestinal epithelial cells (sIECs) play a key role in CeD. These cells are major sources of immunomodulatory factors in the intestinal micromilieu through which they interact with and profoundly influence immune cells and their response to gluten. For example, a previous study has shown that the macrophage response to gliadin is influenced by sIECs (5). Additionally, evidence suggests that sIECs may directly contribute to the inflammatory cascade that characterizes CeD through the production of pro-inflammatory cytokines. For example, previous studies have reported that sIECs can produce the pro-inflammatory cytokines IL-15 and IL-8 in the presence of gliadin, which can activate and recruit immune cells to the intestinal mucosa (2). it is also well known that active CeD causes poorer absorption of nutrients in both children and adults due to this chronic inflammation and intestinal damage (6, 7). This can lead to not only malnutrition, but also development and growth issues in children. After switching to a gluten-free diet and allowing sIECs to heal, normal levels of nutrient absorption can be restored and inflammation can subside, along with many other symptoms that are associated with CeD (8).

Although poor nutrient digestion and malabsorption are well-documented symptoms of active CeD, there are critical gaps in our knowledge regarding the specific dysregulated metabolic processes in sIECs underlying these symptoms. In fact, alterations in sIEC metabolism not only affect nutritional absorption, but also other aspects of CeD such as the intestinal microenvironment, intestinal barrier integrity, immune cell interactions, and the overall inflammatory response to gluten. Addressing this knowledge gap can help unravel the intricate interplay between genetics, metabolism, immunity, and environmental factors that underlies the CeD pathogenesis. Therefore, studying the metabolism of sIECs offers promising potential for the development of new treatment strategies.

GEnome-scale Models (GEMs) of metabolism are an ideal tool for studying sIEC metabolism. These models encapsulate all metabolic reactions encoded by metabolic genes in the genome of an organism and can be constructed from sequenced and annotated genomes. A reconstructed GEM can be also contextualized using transcriptional data. For human cells, one can use transcriptional profiles from specific cell types to infer enzymes or reactions that are active in a cell or tissue and construct cell- or tissue-specific GEMs. The reconstructed GEMs can then be computationally simulated using Constraint-Based Reconstruction and Analysis (COBRA) methods to predict various system-level metabolic properties of the organism such as the growth capacity, nutrient absorption, metabolite secretion, and internal reaction fluxes (9).

In this study, we aimed to investigate the metabolic landscape of sIECs in CeD by utilizing GEMs of metabolism. To this end, we constructed patient-specific GEMs of sIEC metabolism for individuals with active CeD, CeD in remission, and non-CeD controls using transcriptional data. Utilizing these GEMs allowed us to gain a deeper understanding of the molecular mechanisms underlying the metabolic alterations in CeD. Several essential metabolic tasks were identified to have altered activity in active CeD and remission CeD compared to non-CeD controls. Additionally, altered sIEC secretion profiles of several metabolites into the bloodstream or intestinal lumen were identified across the study groups. We further utilized the identified differentially active metabolic tasks to propose potential drugs that have the potential to bring back these altered metabolic functionalities to normal levels in CeD patients.

## Results

### Reconstruction of personalized sIEC GEMs

Patient-specific GEMs of sIEC metabolism were created computationally using a previously published sIEC GEM (10) as a basis and patient-derived transcriptional data (11). This base sIEC GEM was constructed based on extensive literature reviews (10). RNA sequencing data collected from the duodenal biopsies of 42 subjects, including patients with active CeD (n = 12), CeD in remission (on a gluten-free diet; n = 15), and healthy (non-CeD) controls (n = 15) were obtained from a previous study (11). To computationally tailor the GEM for each individual in the cohort, each patient’s transcriptional profile was incorporated into a constraint-based model of the base sIEC GEM using a new method (**Figure 1A** and **Methods**). Constraint-based methods allows for the computation of reaction fluxes within a GEM of metabolism, offering insights into the complex interplay between numerous metabolic reactions and pathways at genome scale. This approach can predict systems-level metabolic properties of sIECs that are not discernible through merely analyzing the transcription profiles of individual genes.

**Figure 1.**
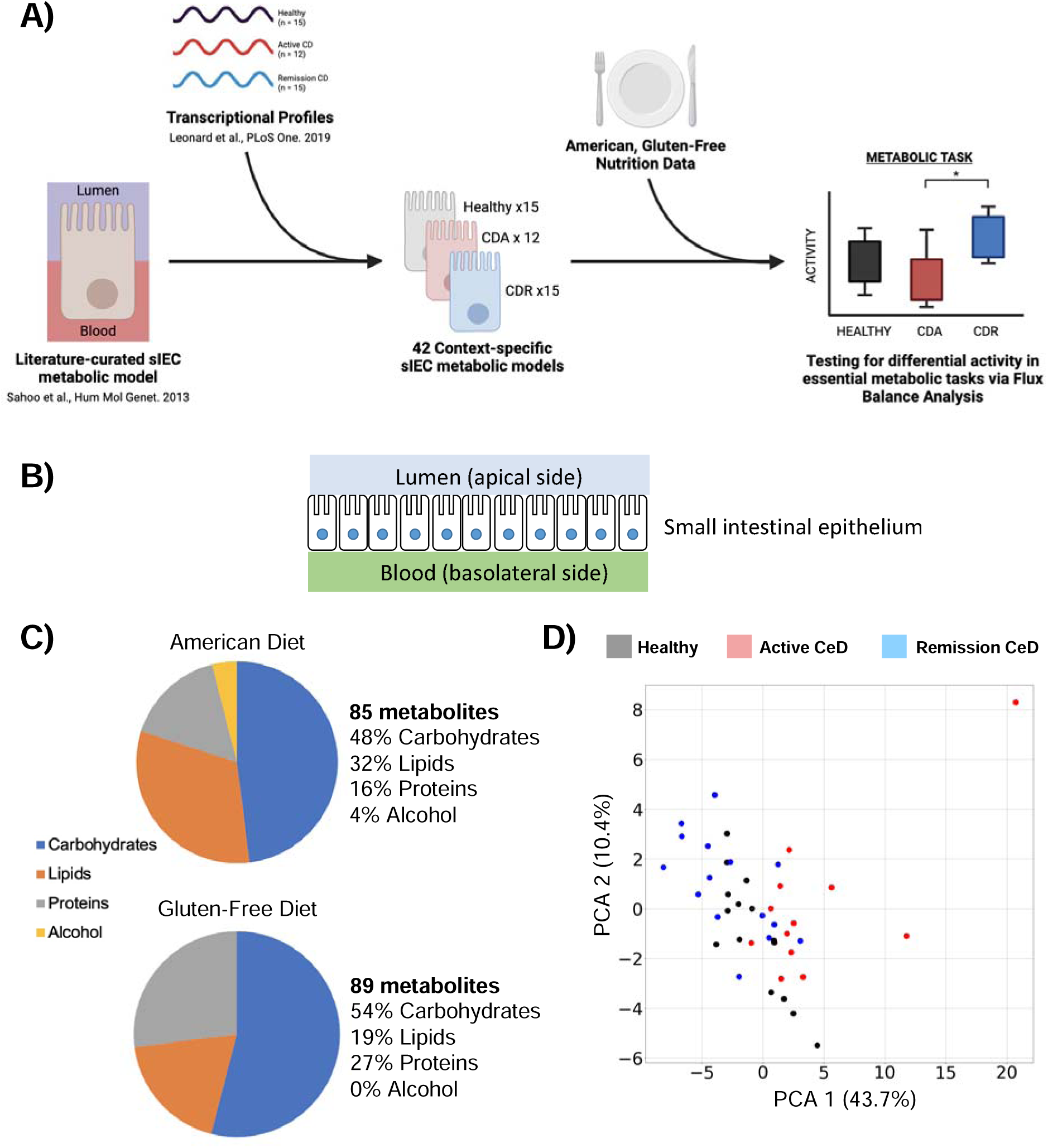
Personalized modeling of sIEC metabolism in CeD using genome-scale models. (A) Schematic representing the construction and simulation of patient-specific sIEC GEMs using transcriptional data. (B) The two extracellular compartments (bloodstream and intestinal lumen) included in sIEC GEMs. (C) Comparison of macronutrient composition for the average American and gluten-free diets used for the simulation of personalized GEMs. (D) Principal Component Analysis (PCA) of sIEC metabolism using the 59 essential metabolic tasks as features.

The personalization of each GEM using the new constraint-based method was achieved by constraining metabolic reactions’ fluxes in the base sIEC GEM as guided by gene expression levels for each individual (see **Methods** and **Supplementary Text**). To further personalize our computational simulations beyond integration with patient-specific gene expression data, we utilized sex-specific information for each individual to parameterize our simulations. To this end, we constrained the growth rate of each sIEC model (the biomass reaction flux within the sIEC GEMs) using values obtained from male and female whole-body models of metabolism developed before (12) (see **Methods**).

The resulting 42 computational sIEC GEMs thus represent the personalized metabolic landscape of sIECs in active CeD, remission CeD, and non-CeD individuals. Each GEM contains two extracellular compartments representing the apical side (intestinal lumen) and the basolateral side (arterial blood or lamina propria). This allows for the simulation of nutrient absorption from the lumen (apical uptake from dietary inputs) and from arterial blood (basolateral uptake), as well as metabolite secretions into the lumen (feces) and blood (**Figure 1B**).

### Computational investigation of sIEC metabolism using personalized GEMs

The personalized sIEC GEMs were simulated under a relevant diet for each group of subjects. An average American diet was formulated and used for the active CeD patients and controls, and an average gluten-free diet was designed and utilized for those with CeD in remission (**Figure 1C**) (10, 13, 14). The nutritional profiles of these two diets exhibit marked differences in their macronutrient and micronutrient compositions. In particular, the Average American diet is characterized by a higher proportion of lipids and alcohol, alongside lower quantities of carbohydrates and proteins (**Figure 1C)**. The detailed micronutrient composition of these diets is provided in **Supplementary Table 1**.

To elucidate metabolic alterations in sIECs associated with CeD, we computationally assessed the capability of sIECs in these subjects to perform 59 defined metabolic tasks essential for growth and functionality, curated previously by Sahoo et al (10) (**Supplementary Table 2**). These tasks encompass a wide range of essential functions examples of which include the synthesis of amino acids and nucleotides, ATP generation via the TCA cycle, the urea cycle, heme synthesis and degradation, and glucose metabolism. Given the fundamental nature of these tasks to sIEC operation and function, they should all be active in a healthy sIEC GEM. In our analyses, we maximized the flux through metabolic reactions representing each task’s activity for each GEM. We further investigated the potential of sIECs in secreting metabolites into the bloodstream and intestinal lumen by maximizing the flux of exchange reactions corresponding to each metabolite. The predicted activity of each metabolic task and the secretion level of each metabolite was recorded and compared between healthy controls, active CeD, and remission CeD groups.

### Overall metabolic landscape of sIECs with respect to essential metabolic tasks across conditions

To gain insights into how the overall metabolic profiles of sIECs with respect to the essential metabolic tasks might be different across the three study groups, we employed Principal Component Analysis (PCA). This analysis shows a partial stratification among the three study groups based on the metabolic profiles of sIECs relative to the essential metabolic tasks **(Figure 1D)**. Subjects in the remission CeD group tend to cluster on the left along the first principal component (PC1), which explains 43.7% of the variance, while subjects with active CeD cluster more toward the right along PC1. This separation implies a divergence in the metabolic capabilities of these two groups. Healthy subjects occupy an intermediate position between the remission CeD and active CeD along PC1. The healthy group shows noticeable overlap with the remission CeD group, demonstrating shared metabolic profiles, and less overlap with the active CeD group, suggesting a metabolic shift in the active CeD group. Less distinct separation between the groups is observed along PC2, which accounts for an additional 10.4% of variance.

Interestingly, an individual from the active CeD group is distinctly positioned away from the main cluster, situated towards the upper right corner. Clinical metadata review revealed no anomalies for this subject, suggesting that its distinctive positioning might reflect unique gene expression profiles characteristic of this subject. Further analysis involving generating a PCA plot generated standardized data (**Supplementary Figure 1, Supplementary Table 3**) confirmed that this data point is an outlier. However, given the lack of any identifiable errors in the collection or handling of biospecimens or data collection for this participant, and considering the potential biological relevance of the atypical gene expression profiles observed, we opted to retain this subject in all subsequent analyses. This allows us to account for the full spectrum of gene expression variability within the active CeD group.

### Essential metabolic tasks with differential activity across conditions

By investigating the activity of the 59 metabolic tasks essential for the functionality of sIECs, a total of 25 metabolic tasks were identified to have significant differential activity between at least one pair of these three groups (Mann-Whitney U, adjusted p < 0.2) (**Figure 2, Supplementary Table 4**). Notably, all these 25 tasks exhibited differential activity between the active CeD and remission CeD groups. Additionally, 15 tasks showed significant differential activity between the active CeD patients and healthy controls while six tasks (L-alanine transaminase, L-alanine secretion, L-arginine secretion, L-proline secretion, 5 10 methylenetetrahydrofolate reductase NADPH, and thymidylate synthase) were differentially active between the remission CeD subjects and healthy controls (**Figure 2**). These metabolic tasks play pivotal roles in maintaining the structural and functional integrity of small intestine, encompassing various aspects such as the gut barrier function, inflammatory response, and oxidative stress regulation. In the subsequent sections, we explore each of these metabolic tasks in greater detail.

**Figure 2.**
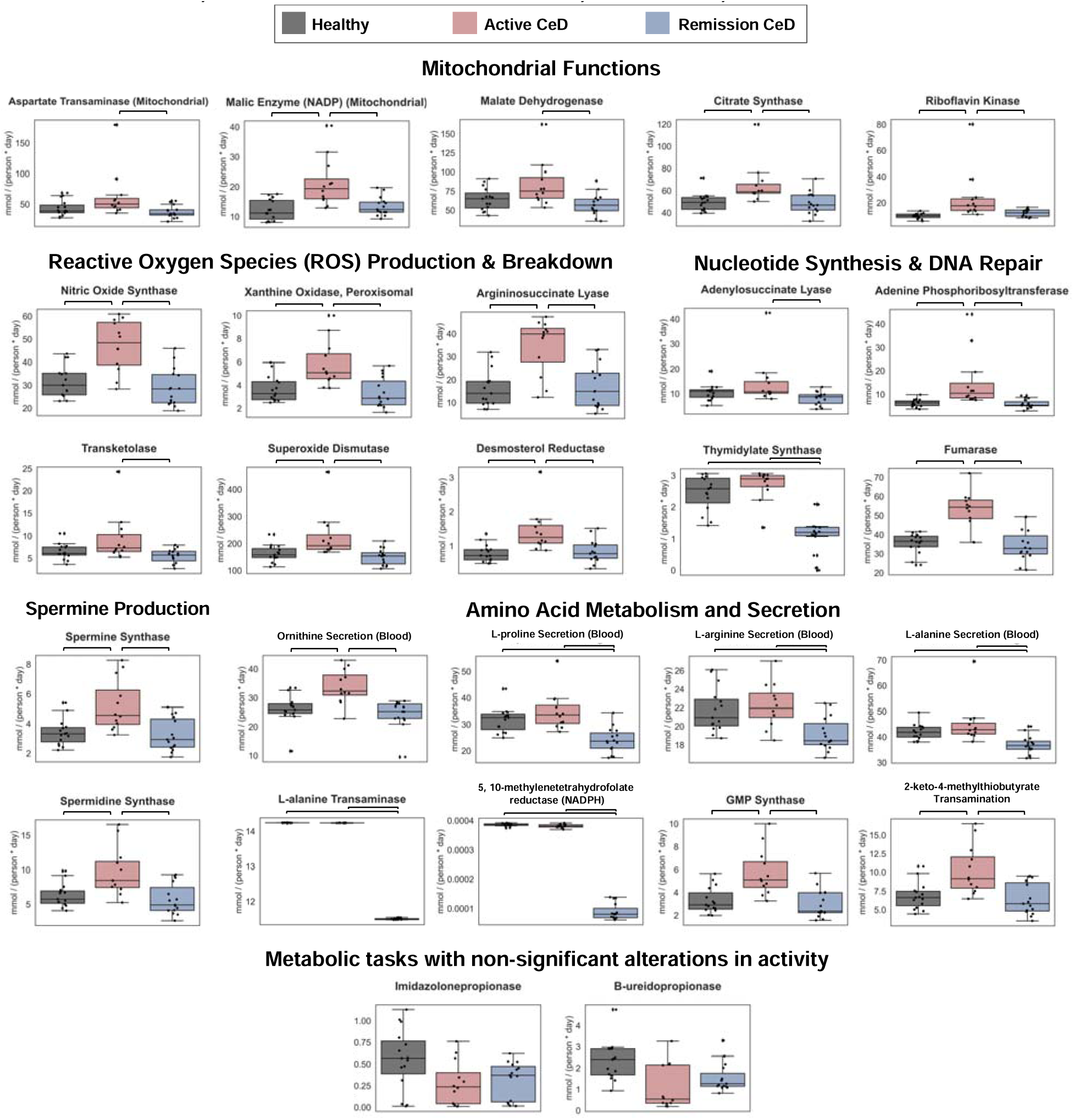
Metabolic tasks in sIECs with significant differential activity across the study groups. The identified 25 metabolic tasks with differential activity across active CeD, remission CeD, and non-CeD controls are involved in various aspects of small intestine metabolism and function. The full list of metabolic tasks tested can be found in **Supplementary Table 2.**

#### Altered mitochondrial functions activity in active CeD

Four metabolic tasks that exhibited differential activity between conditions were directly or indirectly related to the TCA (tricarboxylic or citric acid) cycle, which occurs in the mitochondria and is responsible for deriving energy from nutrients. These tasks include mitochondrial aspartate transaminase, mitochondrial malic enzyme, malate dehydrogenase, citrate synthase, and riboflavin kinase all of which showed elevated flux in active CeD (**Figure 2**).

Mitochondrial aspartate transaminase catalyzes the synthesis of glutamate and oxaloacetate (an important intermediates in TCA cycle). The serum concentration of this enzyme is a commonly used marker for liver health (15). This enzyme displayed significant increased activity in active CeD compared to remission CeD. Elevated aspartate transaminase blood levels have been documented in treatment-naive children and adults with CeD (16, 17). Additionally, transaminase activity was reported to return to normal levels after following a gluten-free diet (17, 18). Interestingly, the co-occurrence of CeD with autoimmune liver diseases has been documented before, strengthening the possibility of shared inflammatory pathways between the two (19).

Mitochondrial malic enzyme, malate dehydrogenase, and citrate synthase also show increased flux in active CeD relative to remission CeD. All three of these enzymes support the production of ATP via the TCA cycle: citrate synthase catalyzes the first step of the TCA cycle, converting acetyl-CoA and oxaloacetate into citrate. Malic enzyme converts L-malate to pyruvate (20) while malate dehydrogenase catalyzes L-malate to oxaloacetate conversion. These three enzymes have varying effects on intestinal inflammation. For example, malate dehydrogenase has been reported to heighten the inflammatory response in patients with Inflammatory Bowel Disease (IBD), and the synthesis of citrate to aid in the production of pro-inflammatory molecules that activate macrophages and dendritic cells. (21, 22). Conversely, malic enzyme promotes the proliferation of sIECs and contributes to barrier maintenance (23).

Riboflavin kinase is another enzyme influencing mitochondrial functions with heightened activity in active CeD. This enzyme is crucial for the metabolism of riboflavin (vitamin B2) to form flavin mononucleotide (FMN), the main precursor for flavin adenine dinucleotide (FAD) synthesis. FAD serves as an essential cofactor in the TCA cycle, and along with FMN, plays a vital role in the electron transport chain by serving as electron carriers (24). Additionally, riboflavin itself is known to have antioxidant and anti-inflammatory properties (25). The elevated activity of malic enzyme and riboflavin kinase in active CeD patients could be a mechanism to compensate for the energy deficit caused by the malabsorption of nutrients due to intestinal damage.

Notably, no genes corresponding to the four metabolic tasks discussed above show significant differences in expression among the groups (Mann Whitney U test, adjusted p < 0.2) (**Supplementary Figure 2**). Aspartate transaminase and citrate synthase are encoded by *GOTL1* and *CS* respectively, while malic enzyme and malate dehydrogenase each is encoded by two genes (*ME2*/*ME3* and *MDH/MDH1B,* respectively).

#### Differentially active metabolic tasks involved in reactive oxygen species (ROS) synthesis and metabolism

Six differentially active metabolic tasks contribute directly or indirectly to ROS production or metabolism in the gut. These include nitric oxide synthase, xanthine oxidase, argininosuccinate lyase, superoxide dismutase, transketolase 2, and desmosterol reductase. These tasks exhibit heightened activity in active CeD compared to the remission CeD and/or control groups (**Figure 2**). Again, no significant changes in the expression levels of the genes producing these enzymes—*SOD2* for superoxide dismutase, *XDH* for xanthine oxidase, TKT, TKTL1, and *TKTL2* for transketolase, *ASL* for argininosuccinate lyase, and *NOS1*/*NOS2*/*NOS3* for nitric oxide synthase—were observed (**Supplementary Figure 2**). ROS can cause oxidative damage and apoptosis in the intestine, compromising the gut barrier integrity. They also impact the immune response: ROS can alter innate immunity by activating macrophages and other innate immune cells towards pro-inflammatory states and have been implicated in many diseases involving chronic inflammation (26). High levels of ROS are a biomarker of CeD and play a role in its pathogenesis by contributing to villi damage (27).

Nitric oxide synthase produces nitric oxide (NO), a compound which plays an important role in regulating and maintaining intestinal barrier integrity; however, it can also cause oxidative stress and damage if present at high concentrations (28). Xanthine oxidase is also involved in the production of ROS, which can polarize immune cells into a pro-inflammatory phenotype (29). Both nitric oxide synthase activity and nitric oxide levels are reported to increase in CeD patients in response to gluten but decrease upon adopting a gluten free diet (30, 31). Argininosuccinate lyase catalyzes the cleavage of argininosuccinic acid to generate arginine, a substrate for nitric oxide synthase. This therefore leads to increased nitric oxide generation, oxidative damage, and inflammation in enterocytes (32). However, increased arginosuccinate lyase activity is also associated with improved epithelial integrity and was reported to improve colitis symptoms in vitro (33) and to prevent necrotizing enterocolitis by decreasing apoptosis and immune cell activation (34).

Unlike nitric oxide synthase, xanthine oxidase, and argininosuccinate lyase, which contribute to ROS synthesis, superoxide dismutase, transketolase 2, and desmosterol reductase are involved in ROS breakdown and regulation. Superoxide dismutase is a main ROS-scavenging enzyme that converts superoxide radicals to hydrogen peroxide, decreasing oxidative stress (35). Transketolase 2 is another enzyme that indirectly contributes to ROS breakdown and regulation: it is involved in the Pentose Phosphate Pathway, which is crucial in the production of NADPH, a co-factor essential for maintaining redox balance. Transketolase 2 has been shown to minimize ROS infiltration into the intestine and support intestinal barrier maintenance in animal models of IBD (36). Desmosterol reductase, which produces cholesterol from desmosterol, is a key enzyme in the cholesterol synthesis pathway (37) that has cytoprotective effects, preventing apoptosis and improving cell survival through the mitigation of ROS production (38).

While counterintuitive, the increased activity of these three ROS-modulating enzymes in active CeD aligns with prior studies. For example, superoxide dismutase elevated activity has been documented in active CeD patients relative to remission CeD and healthy controls (39). It was proposed that gliadin may cause an imbalance in antioxidant activity, leading to the increased odds of lesions in the intestinal mucosa (40). The increased activity of these three enzymes may also represent the body’s response to counteract the enhanced activity of nitric oxide synthase and xanthine oxidase in active CeD patients.

#### Differentially active metabolic tasks involved in nucleotide synthesis and DNA repair

Four functions with variations in activity across the study groups engage with nucleotide synthesis and DNA repair essential for sIEC growth and regeneration. These include adenylsuccinate lyase, adenine phosphoribosyltransferase, thymidylate synthase, and fumarase. Adenylsuccinate lyase and adenine phosphoribosyltransferase participate in purine biosynthesis and purine salvage pathways, respectively (41, 42). Both of these show a significantly higher activity in the active CeD group relative to remission CeD (**Figure 2**). Thymidylate synthase, which exhibits elevated activity in the active CeD and healthy controls relative to remission CeD, also participates in DNA synthesis as a key regulator of thymidine (one of the nucleotides in DNA) (43). Related to these functions is cytosolic fumarase, which catalyzes the formation of L-malate from fumarate, and its activity contributes to the DNA damage response (44). Fumarase shows heightened reaction fluxes in the active CeD group compared to the other two groups.

The enhanced activity of these functions in active CeD may be due to the increased damage of cells in active CeD, leading to the activation of pathways that contribute to DNA repair and synthesis. The diminished activity of thymidylate synthase in remission CeD compared to healthy controls might reflect the reduced need for DNA synthesis and repair due to the healing of the intestine after following a gluten-free diet or due to the nutritional differences between the two.

#### Elevated spermine synthesis in CeD

Spermine synthase and spermidine synthase, two enzymes responsible for spermine synthesis, showed significantly higher activity in the active CeD patients compared to both healthy controls and remission CeD groups (**Figure 2**). Spermine, the product of spermine synthase, is a polyamine that promotes cellular growth, modulates gut epithelium integrity, and supports healthy epithelial barrier function (45). In CeD, spermine has been shown to competitively inhibit the tTG enzyme. tTG inhibitors are considered as potential therapeutics for CeD (46). Spermine has also been characterized as an inhibitor of inflammation through protecting against oxidative stress and inhibiting inflammatory cytokine synthesis within the innate immune system (47). Spermidine, the product of spermidine synthase, is a precursor for spermine biosynthesis and has also been shown to attenuate gliadin’s toxic effects on sIECs (48), induce CD4^+^ T cell differentiation towards regulatory T cells (Treg) suppressing pro-inflammatory responses, and polarize macrophages towards an anti-inflammatory phenotype (47, 49).

These reports imply a protective effect for spermine synthase and spermidine synthase on the gut barrier function and intestinal inflammation, yet we observe a higher activity for them in active CeD patients. While counterintuitive, some of these observations are indeed consistent with existing literature for other inflammatory diseases. For example, spermine, along with other polyamines, were reported to be elevated in the blood serum and colonic mucosa of colorectal cancer patients (50) and in patients with acute colitis (49, 51). Interestingly, spermidine is also currently being explored as a therapy to inhibit tTG activity, inhibiting inflammation due to gliadin deamidation (52). The higher activity of these two metabolic functions in active CeD patients in our results and in prior studies may represent a compensatory response by the body to mitigate the inflammation and damage to the intestine in CeD.

#### Amino acid production and secretion is altered in both active and remission CeD patients

We observed significant altered activity in eight metabolic tasks related to amino acid metabolism and transport in the active and remission CeD groups. Four of these tasks engaged in the secretion of amino acids into blood. Specifically, L-alanine, L-arginine, and L-proline exhibited elevated secretion flux into blood in the active CeD and healthy groups relative to the remission CeD group. Additionally, ornithine showed a significant increased secretion in the active CeD group compared to both remission CeD and controls **(Figure 2**). The observed elevated secretion of these amino acids in active CeD relative to remission CeD is in line with previous studies reporting elevated amino acid levels in blood plasma collected from individuals with active CeD (53, 54).

The other four tasks with altered activity are related to amino acid metabolism and include L-alanine transaminase, 5, 10-methylenetetrahydrofolate reductase (NADPH), guanosine monophosphate (GMP) synthase, and 2-keto-4-methylthiobutyrate transamination. Two of these tasks, namely L-alanine transaminase and 5, 10-methylenetetrahydrofolate reductase (NADPH), exhibit significantly reduced activity in remission CeD compared to active CeD and controls. L-alanine transaminase is responsible for the intestinal catabolism of dietary compounds to synthesize alanine, an important nitrogen source for extraintestinal tissues (53). The increased activity of alanine transaminase in children and adults with active CeD has been documented before (16). 5, 10-methylenetetrahydrofolate reductase (NADPH) is an enzyme implicated in folate cycle and is a key contributor to homocysteine (an amino acid) production. Prior research shows elevated homocysteine levels in CeD patients (55), which is inconsistent with our findings showing no significant difference between active CeD and controls. The diminished activity of these two functions in remission CeD relative to the other two groups might be due to the differences in the gluten-containing versus gluten-free diets in these groups.

GMP synthase and 2-keto-4-methylthiobutyrate transamination are related to glutamate (glutamic acid) production and also showed higher fluxes in the active CeD patients compared to remission CeD and/or healthy controls (**Figure 2**). Glutamate has been reported to be elevated in the plasma of active CeD patients (53). It has been also shown to enhance intestinal barrier integrity and antioxidant functions, and to help reduce injury to the intestine caused by inflammation (56).

CeD patients have been shown to have impaired capacity for absorbing amino acids and peptides compared to healthy individuals due to the damage to intestine (57), potentially contributing to increased activity of enzymes involved in amino acids synthesis. Furthermore, it has been suggested that the elevated activity and plasma levels of amino acids in active CeD patients may contribute to the systemic inflammation seen in CeD (53).

#### Non-significant alterations in essential metabolic tasks

In addition to the 25 metabolic tasks with significant differential activity discussed above, we observed marked, albeit non-significant, differences in the activity of an additional 24 metabolic tasks as shown in **Supplementary Figure 3**. These included two metabolic tasks, namely imidazolone propionase and B-ureidopropionase, that had a reduced activity in active CeD relative to the other two groups (**Figure 2**).

### Metabolites differentially secreted into the bloodstream and intestinal lumen

To investigate the capacity of sIECs in secreting different metabolites into the blood circulation and intestinal lumen across CeD conditions, the maximum secretion potential of various metabolites was quantified. These metabolites correspond to 166 exchange reactions within the sIEC GEMs that transport metabolites from the cytosol into either the bloodstream or lumen.

Through this analysis, we identified 55 distinct metabolites (other than those already discussed as essential metabolic tasks) with significant variations in secretion flux into the blood or lumen between at least two conditions (Mann-Whitney U, adjusted p < 0.2, **Figure 3** and **Supplementary Table 5**). Of these, 53 showed differential secretion into the bloodstream, one (nitric oxide) into the lumen, and one (hydrogen) into both the blood and lumen across conditions. These metabolites span various categories such as amino acids, vitamins, antioxidants, and fatty acids, among others.

**Figure 3.**
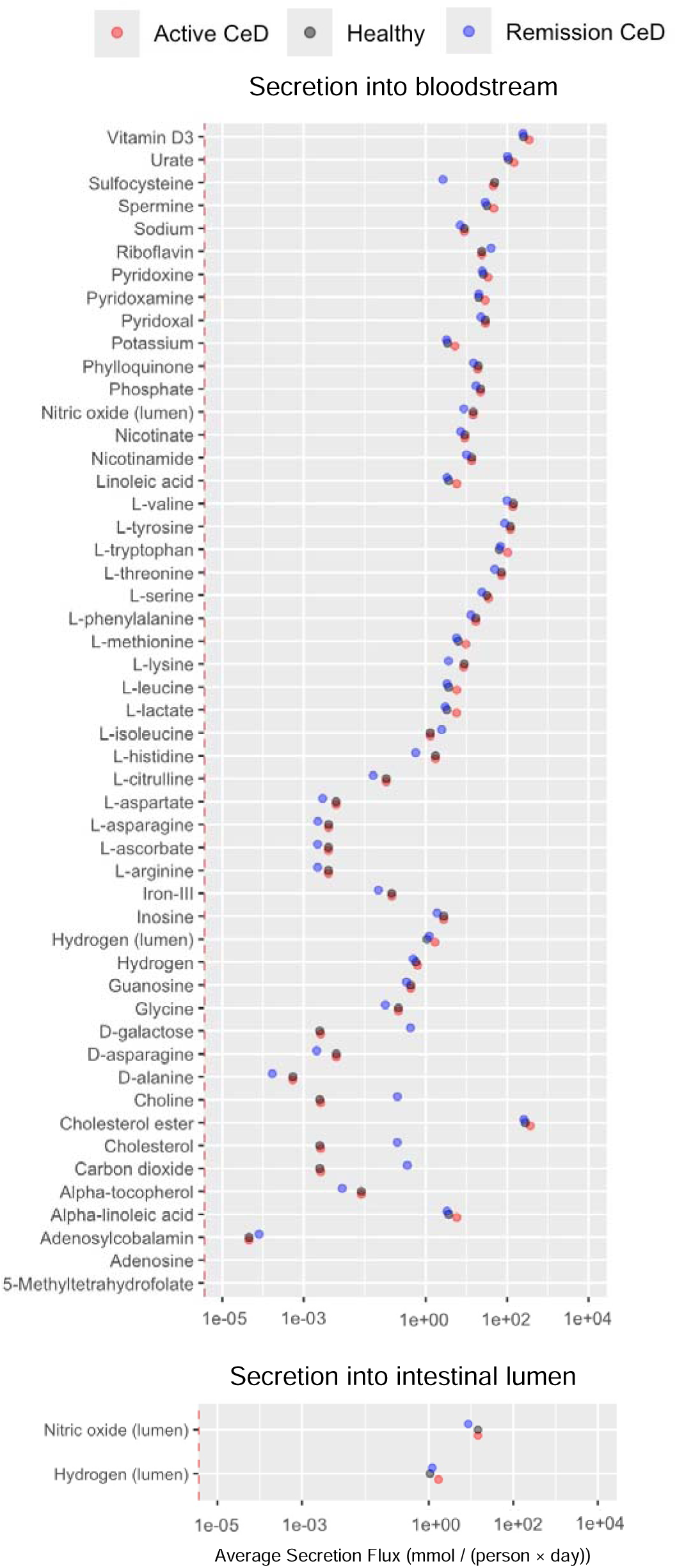
Metabolites with differential secretion profiles into the bloodstream and intestinal lumen across study groups. Metabolites listed have significant differential secretion between at least one pair of conditions (Mann-Whitney U Test, p < 0.2). Error bars represent the standard error of each secretion level. A full list of metabolites with differential secretion profiles can be found in **Supplementary Table 5**.

Notably, 20 of these metabolites are various isoforms of essential and non-essential amino acids. Of these, 16 amino acids had higher secretion fluxes into the blood in active CeD and healthy controls relative to remission CeD. Additionally, L-methionine, L-tryptophan, and L-leucine exhibited increased secretion into blood in active CeD relative to the other two conditions, while L-isoleucine showed increased secretion in remission CeD compared to active CeD and healthy groups. These altered amino acid secretion profiles reflect changes in the metabolic state of sIECs in both active and remission CeD.

Among other metabolites with differential secretion profiles are those related to ROS. Specifically, nitric oxide secretion into the lumen was higher in the active CeD, and healthy groups compared to remission CeD, and urate secretion into the blood was higher in active CeD compared to both healthy controls and remission CeD. As noted before, nitric oxide is an ROS that contributes to inflammation when present at high concentrations and its increased secretion levels in active CeD is consistent with the elevated activity of the metabolic task nitric oxide synthase in the active CeD patients (**Figure 2**). Conversely, urate is an antioxidant reported to have higher serum concentration in adults with CeD, which aligns with our findings (58).

Another metabolite of interest is linoleic acid and its isoform alpha-linoleic acid, which both exhibit higher secretion into the bloodstream in active CeD compared to remission CeD and healthy controls. Linoleic acid is an essential fatty acid and its derivatives have pro-inflammatory properties (59). Previous studies have reported elevated serum levels of unsaturated fatty acids, including linoleic acid, in patients with active CeD (60, 61). High levels of linoleic acid have also been associated with a higher risk of colorectal cancer (62, 63).

Differential secretion of other micronutrients was also observed. For example, we recorded significantly reduced secretion into the blood for B vitamins adenosylcobalamin and 5-methyltetrahydrofolate in active CeD relative to remission CeD. This is consistent with micronutrient deficiencies often observed in CeD patients (64) due intestinal damage. Our results also show increased secretion levels of other vitamins in active CeD compared to remission CeD or controls examples of which include vitamin D3, pyridoxamine, and pyridoxine. Additionally, vitamin E (alpha tocopherol), vitamin K (phylloquinone), vitamin C (L-ascorbate), and B vitamins pyridoxal, nicotinamide, and nicotinate exhibited diminished secretion in remission CeD relative to the other two groups, likely due to nutritional differences between these groups.

### Probing sIEC metabolic adaptability for performing essential functions

A shadow price analysis was conducted for each personalized GEM to evaluate sIEC metabolic adaptability to perturbations for performing the 59 essential metabolic tasks and metabolite secretions. This analysis did not identify a significant difference in the overall sIEC adaptability for performing these functions (the sum of non-zero shadow prices across all tasks or metabolites) between the study groups (Mann-Whitney U, p < 0.2; **Supplementary Table 6**). However, when assessing the adaptability for individual metabolic tasks or metabolite secretions, significant differences between the groups were observed for two tasks: ornithine secretion into the bloodstream and carboxylic acid dissociation (**Figure 4**). The non-zero shadow prices count for ornithine secretion was significantly lower in the active CeD patients, indicating a lower adaptability, compared to the remission CeD. Interestingly, this is despite the elevated secretion of ornithine into blood observed in the active CeD patients **(Figure 2).** Conversely, the adaptability for carboxylic acid dissociation, as reflected by non-zero shadow prices count, was significantly diminished in both active and remission CeD groups relative to healthy controls. This function is responsible for the production of carbonic acid, which then dissociates into HCO_3_^-^ ions. HCO_3_ ions neutralize acids like gastric hydrochloric acid that may harm the gastrointestinal tract. These findings indicate a lower flexibility of active CeD and remission CeD patients to perform metabolic processes related to these two functions.

**Figure 4.**
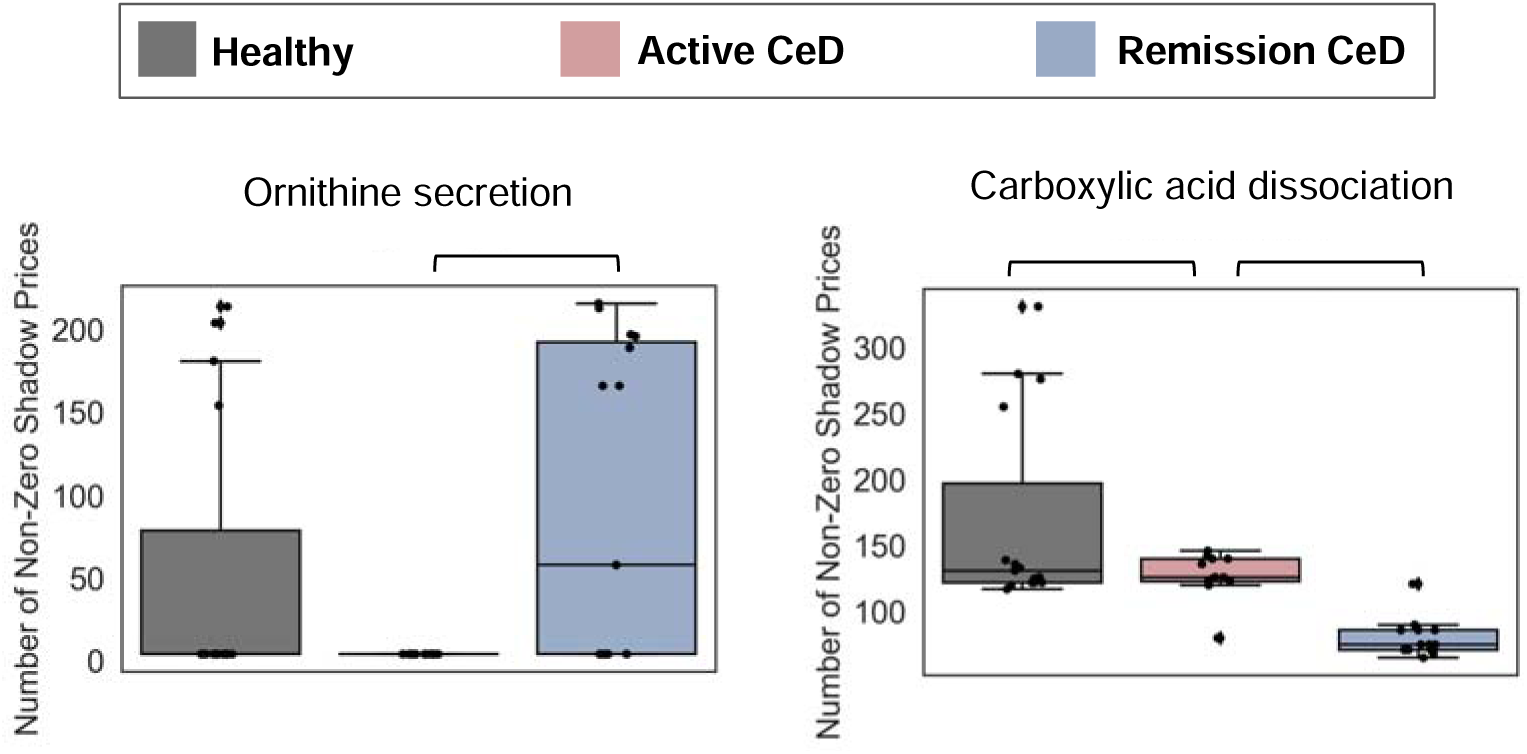
Analysis of sIEC metabolic adaptability across conditions. In this analysis, a higher count of non-zero shadow prices represent an increased flexibility of the metabolic network for handling perturbations. Metabolic tasks (ornithine secretion, carboxylic acid dissociation) with significant differential activity across the groups are shown.

### Sensitivity analysis

To evaluate if the identified outlier from the active CeD group (Figure 1D) may distort our statistical analysis of metabolic tasks, metabolite secretions, and sIEC metabolic adaptability, we conducted a sensitivity analysis excluding this subject. This exclusion rendered only three metabolic tasks—transketolase, adenylsuccinate Lyase, and spermine synthase—statistically non-significant, although the overall pattern of differential activity for these tasks remained unchanged.

### Drug target analysis

We next explored potential drugs that can restore the altered activity of the differentially active metabolic tasks in active CeD or remission CeD patients to those of healthy ones. Out of the 25 differentially active metabolic tasks, we focused our attention on six metabolic tasks whose observed altered activity had detrimental effects on the intestine and contribute to CeD development. These tasks include citrate synthase, aspartate transaminase, and malate dehydrogenase (promoting pro-inflammatory immune cell activation), as well as nitric oxide synthase, xanthine oxidase, and argininosuccinate lyase (implicated in ROS synthesis) (see the previous sections for details). Gene-protein-reaction rules in GEMs were assessed to determine the gene or genes that are responsible for each of these tasks. This resulted in 6 genes (MDH, NOS1, NOS2, NOS3, CS, ASL) that serve as candidate drug targets.

We additionally conducted Flux Coupling Analysis (FCA) (65) to pinpoint reactions in the GEMs that are fully coupled the specific reactions representing these six metabolic tasks. Modulating the expression of the genes encoding reactions that are fully coupled with the target reactions for a metabolic task will also impact the activity of that metabolic task the same way. This analysis revealed four additional reactions that are fully coupled with the six metabolic tasks. These coupled reactions were linked to two additional genes (SLC25A12, SLC25A13, ASS1) yielding a total of 10 genes to target.

Next, we queried the DrugBank database to identify potential drugs–including both FDA-approved and those in clinical trials—that target these 10 genes by up- or down-regulating their expression levels. With this analysis, we identified 22 drugs that target the expression of at least one gene responsible for the six differentially active metabolic tasks or reactions fully coupled to them (Figure 5, **Supplementary Table 7**). Of note, five of these drugs (acetaminophen, cisplatin, cyclosporine, silicon dioxide, valproic acid) target multiple genes.

**Figure 5.**
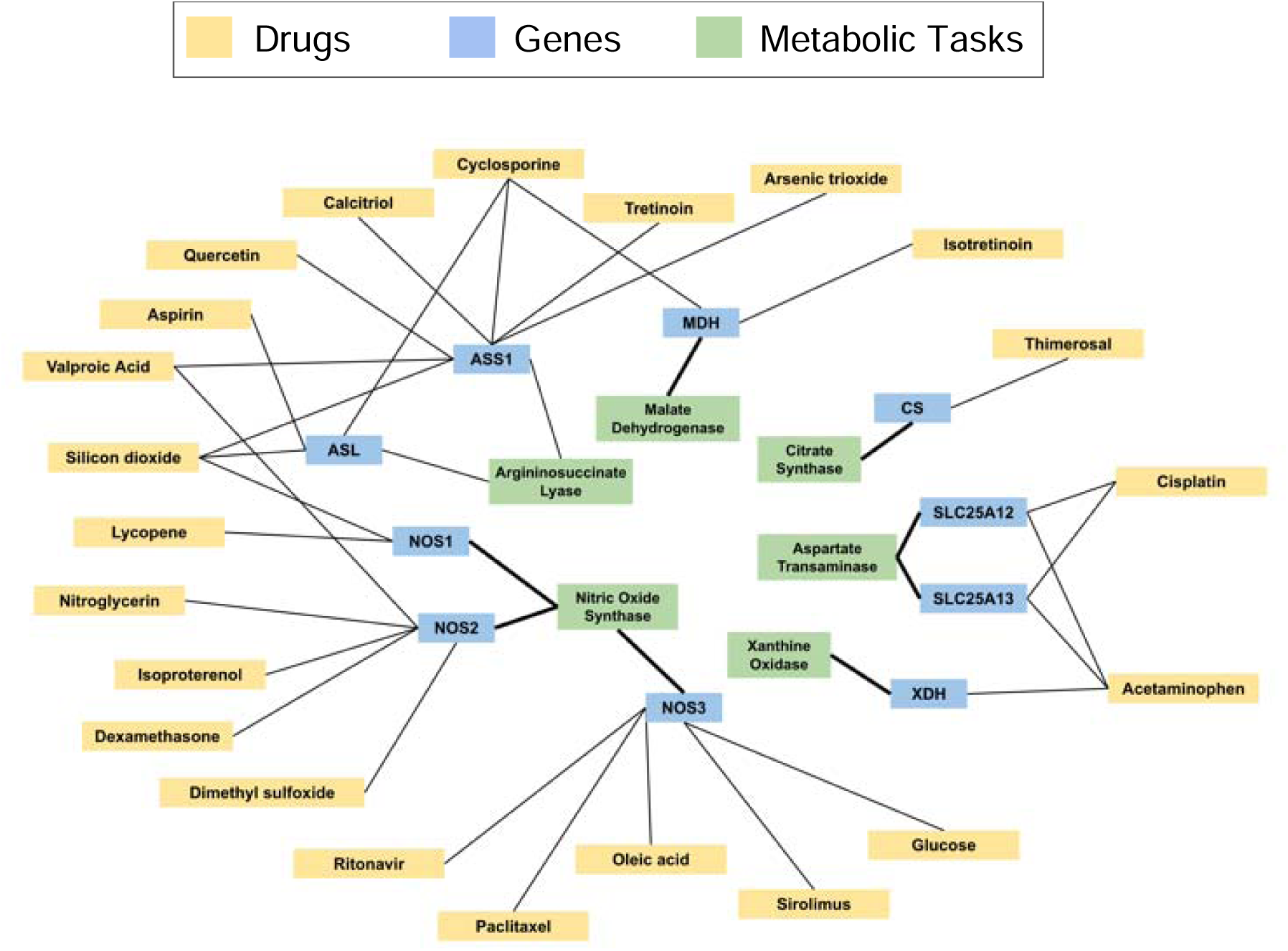
Drugs targeting the genes for altered metabolic tasks in sIECs in CeD. Existing FDA-approved drugs targeting 10 genes encoding six differentially active metabolic tasks with adverse effects on sIECs in CeD are shown. Lines represent gene-to-reaction links or gene-drug interactions. All drugs shown suppress the expression of the genes linked to them. The full list of these drugs can be found in Supplementary Table 7.

## Discussion

In this study, we sought to systematically dissect the metabolic landscape of sIECs in CeD by leveraging genome-scale modeling. To this end, patient-specific GEMs of sIEC metabolism were constructed by incorporating transcriptional profiles from 42 subjects with active CeD, CeD in remission, and healthy controls as well as sex-specific parameters into our computational framework. By simulating an average American diet and a gluten-free diet using these personalized GEMs, we computationally investigated the capability of sIECs in performing 59 curated essential metabolic tasks. Analyzing the overall metabolic state of each GEM with regard to the metabolic tasks using PCA revealed a distinct metabolic state in active CeD patients compared to those with CeD in remission and healthy controls (**Figure 1D).** Additionally, we observed a marked overlap between the clusters for healthy and remission CeD individuals indicating the intestinal healing that occurs within CeD patients after adopting a gluten-free diet. While the remission CeD and active CeD show a more pronounced stratification, there is a slight interspersion between the two (near the center of PC1), which could indicate the potential of a few active CeD patients to go into remission, or the potential for a few remission CeD patients to experience relapse and re-develop CeD symptoms.

We next investigated the activity of each essential metabolic task across the three study groups. This analysis revealed 25 metabolic tasks with significant differential activity between at least one pair of these groups (**Figure 2**). Many of these differentially active metabolic tasks are implicated in important processes involved in CeD pathogenesis such as gut barrier maintenance, immune system modulation, and nutrient absorption and metabolism. A subset of these tasks which exhibited elevated activity in CeD contribute to CeD pathogenesis by inducing a pro-inflammatory response (malate dehydrogenase and citrate synthase) or by compromising the gut barrier via ROS production (nitric oxide synthase, xanthine oxidase, and argininosuccinate lyase). Conversely, we identified several other metabolic tasks with elevated activity in active CeD patients that were known from existing literature to have protective effects. Notably, some of these tasks can promote anti-inflammatory responses and/or improve gut barrier integrity (malic enzyme, riboflavin kinase, superoxide dismutase, transketolase 2, desmosterol reductase, spermine synthase, spermidine synthase, GMP synthase, and 2-keto-4-methylthiobutyrate transamination). This seemingly counterintuitive observation for these metabolic tasks highlights the complex relationships between these critical metabolic functions and the pathophysiology of CeD. A highly likely scenario for the heightened activity of the metabolic tasks with protective effects in the active CeD patients may represent a compensatory response by the body to mitigate intestinal damage due to the immune response to gluten. Other factors that may also contribute to these observations may include the malabsorption of nutrients due to intestinal damage, the disruption of metabolic pathways triggered by disease, or the complex interactions between multiple metabolic processes in the disease state (2). These insights underscore the multifaceted nature of metabolic adaptations in response to the physiological challenges imposed by CeD. We additionally detected (non-significant) alterations in the activity of certain metabolic tasks between individuals with CeD in remission and controls (e.g., argininosuccinate lyase, malic enzyme, imidazolone propionase, and B-ureidopropionase) indicating that while their overall metabolic profiles are similar, there are specific alterations in the metabolism of those with CeD in remission.

Notably, for numerous metabolic functions that displayed significant differential activity, we did not observe any significant changes in the expression levels of the genes encoding them. This highlights the merits of employing genome-scale modeling in elucidating the phenotypic outcomes that arise from the complex interplay among multiple metabolic pathways. This system-wide perspective offers non-intuitive insights that transcend those provided by the isolated examination of expression levels for individual genes. GEMs thus deliver a more holistic understanding of sIEC metabolism and function.

In addition to the essential metabolic tasks, we explored differences in the secretion profiles of various metabolites by sIECs into both the blood and intestinal lumen. This revealed 52 distinct metabolites with significant differential secretion into blood and two into the lumen between at least one pair of study groups. These variations in secretion profiles reflect alterations in sIEC metabolism due to the disease state and/or dietary interventions.

Of note, for a number of metabolic tasks, including L-alanine, L-arginine, and L-proline secretion and L-alanine transaminase, 5 10 methylenetetrahydrofolate reductase NADPH, we recorded significant diminished activity or secretion in remission CeD compared to both controls and active CeD subjects. The same pattern was also observed for several metabolite secretion profiles into the bloodstream spanning L-citrulline, adenosine and guanosine, Vitamin E (alpha tocopherol), vitamin K (phylloquinone), vitamin C (L-ascorbate), as well as B vitamins pyridoxal, nicotinamide, and nicotinate. These observations either have not been documented before or are inconsistent with existing literature reporting a lower activity or secretion in active CeD patients. The reduced activity of these metabolic tasks and metabolites secretion levels in the remission CeD group can be attributed, in part, to the nutritional deficiencies inherent to the gluten-free diet, which could alter metabolic pathways and secretion profiles in these subjects. This phenomenon could also stem from a combination of other inter-connected factors. These may include incomplete intestinal healing in remission CeD patients (where enterocyte function remains suboptimal), a shift in metabolic demands prioritizing tissue repair, persistent residual inflammation influencing metabolic pathways, or the complex adaptive regulatory mechanisms in remission CeD aimed at managing persistent residual inflammation through adjusting metabolic and transporter activities. These elements reflect the complex interplay of nutritional factors, physiological adaptations, and residual effects even in the remission phase of CeD.

While probing sIEC overall metabolic adaptability to perturbations did not reveal a significant difference across the study groups, significant alterations were noted for two individual metabolic tasks. Specially, a significantly lower count of non-zero shadow prices for ornithine secretion was detected in active CeD patients—despite higher ornithine secretion levels in active CeD. This suggests that the metabolic system in the active CeD state is operating under more genetic or epigenetic constraints, due to inflammation or tissue damage, limiting its ability to utilize alternative pathways or adapt to increased metabolic demands in this state.

Following a gluten-free diet is currently the only available treatment for CeD, which is very difficult to undertake as gluten is present in the vast majority of foods, and even trace amounts of gluten can induce damage to the small intestine. Moreover, this treatment is not completely effective as 30% of patients still report ongoing symptoms and 20% have persistent enteropathy despite following a gluten-free diet (8). In this study, we conducted a drug target analysis, which identified 22 FDA-approved drugs that target the genes encoding six metabolic tasks with detrimental activity that adversely affect sIECs proper functioning and contribute to CeD pathogenesis, By restoring the normal expression levels of these target genes, and consequently rectifying the activity of the respective metabolic tasks, these medications offer promising therapeutic options for ameliorating CeD symptoms—especially benefiting those patients that persistently suffer despite adhering to a gluten-free diet. Many of these drugs have immunosuppressive or anti-inflammatory properties. in particular, cyclosporine, dexamethasone, sirolimus, and isotretinoin have been approved by the FDA as anti-inflammatory or immunosuppressant drugs in multiple disease contexts such as rheumatoid arthritis and cancer (66). Additionally, two of these drugs, cyclosporine, and methotrexate, are already used to treat other inflammatory diseases of the gut such as Crohn’s Disease and Ulcerative colitis. However, none of these medications are currently being investigated to treat symptoms of CeD. Other drug groups identified in this analysis include chemotherapy drugs and other anti-tumor agents, corticosteroids, retinoids, as well as common over-the-counter vitamins and supplements. Of these, corticosteroids such as prednisolone have been investigated to potentially aid CeD patients who do not improve upon starting a gluten-free diet; however, the corticosteroid dexamethasone which was identified in this analysis has not yet been studied in the context of CeD (67).

It is worth highlighting some limitations of this study. While GEMs of metabolism can adeptly model metabolic reactions and the interactions between reaction products and metabolic pathways, it is essential to acknowledge their inherent constraints and limitations. Notably, GEMs do not encompass regulatory interactions, allosteric regulation of enzymes, and the effects of cell signaling via cytokine production or alternative mechanisms, among others. Additionally, it is crucial to recognize that GEMs primarily function as tools for hypothesis generation. The predictions derived from these computational models necessitate validation through in vitro or in vivo studies and/or clinical trials. For instance, determining if the drugs identified in this study can effectively modulate or restore healthy sIEC metabolic processes or to potentially alleviate CeD symptoms warrants thorough examination and experimental and clinical validation. For example, we have recently explored the protective role of a strain of *Bacteroides vulgatus* in ameliorating damage to sIECs in CeD by employing gut organoid models (PMID: 38177249). These models provide a valuable foundation for future research to evaluate the efficacy of the drugs identified in the current study.

## Conclusion

Overall, our study paves the way for a rigorous and quantitative exploration of metabolic dysfunction in CeD and other chronic human diseases, through leveraging personalized GEMs of metabolism patient-specific incorporating transcriptional data. As demonstrated, this approach goes beyond classical differential gene expression analysis, allowing for the observation of intricate interactions within the complex network of gene products, metabolites, metabolic reactions and pathways in disease. These findings offer new insights into the dysregulation of specific metabolic processes within sIECs in CeD, shedding light on novel avenues for therapeutic intervention and personalized treatment strategies. Specifically, this research has pinpointed promising FDA-approved drug candidates capable of rectifying the altered metabolic functions contributing to CeD pathogenesis, offering a pathway to repurposing existing drugs for CeD treatment. These findings mark an essential step towards alternative and effective treatment options and improved quality of life for individuals suffering from this disease.

## Methods

### Personalized sIEC GEM modeling

An existing literature curated GEM of metabolism for sIECs was used as a baseline model.(10) This GEM contains 1,282 reactions and 433 unique metabolites located in five intercellular compartments (cytosol, nucleus, mitochondria, peroxisome, and endoplasmic reticulum) and two extracellular compartments representing the intestinal lumen and the bloodstream to reflect the luminal (apical) and basolateral sides of the sIEC. Personalized computational sIEC models were then constructed from this baseline model by incorporating gene expression (RNA sequencing) data collected from the duodenal biopsies of subjects with active CeD, CeD in remission and healthy controls (11) into Flux Balance Analysis (FBA) models. This was done by constraining the reaction fluxes in GEMs according to gene expression data for each subject (McCreery and Zomorrodi, manuscript under preparation). To create these personalized sIEC GEMs, genes were grouped into 20 clusters based on their level of expression across all samples using the StanDep package (68), implemented in MATLAB 2018. Within each of these clusters, gene expression was normalized to ensure uniformity between samples within that cluster. This clustering strategy allows for the retention of genes with low expression levels and safeguards against their removal by applying a universal threshold. Normalized gene expression data are used to impose soft constraints on the lower and upper bound on reaction fluxes in the network. We then minimize deviations from the imposed bounds on reactions in the GEM (i.e., deviations from gene expression data) as the objective function. The detailed mathematical formulation for this problem is provided in the **Supplementary Text**.

**In silico diets:** An average American diet was obtained from food intake data recorded by the 2007-2008 National Center for Health Statistics’ National Health and Nutrition Examination Survey (**69**) **(Supplementary Table 8).** From this survey, a diet was reconstructed by calculating the average intake of each item listed in the survey’s 226 food and drink groups. Similarly, an average gluten-free diet was created from a list of 21 food items within a typical gluten free diet reported by CeD patients at Massachusetts General Hospital, Boston, MA, USA **(Supplementary Table 9)**. This list of food items from each diet was then converted to bounds on uptake fluxes of 85 and 89 metabolites, for the Average American and Glute-free diets, respectively, using the Diet Designer tool in the Virtual Metabolic Human database (14). Aside from these nutrients acquired from the diet, the uptake of oxygen, bicarbonate, and asparagine from the blood compartment, as well as bicarbonate from the lumen compartment, were allowed for both diets as proposed in (10). This ensures flux consistency in the models for both the Average American and Gluten-free diets. For each metabolite in a diet, its corresponding exchange reaction’s lower bound was set to reflect that metabolite’s uptake. The lower bound for exchange reactions responsible for metabolites that were not included in these diets were set to zero.

### Computational simulation of GEMs

To incorporate sex as a variable in our analyses, we constrained each GEM’s biomass reaction between a specific lower and upper bound that were gender specific. The biomass reaction represents the metabolites and resources necessary for cellular maintenance and growth for sIEC. To calculate the gender-specific bounds, we utilized the organ-resolved male and female Whole-Body Model (WBM) of metabolism developed by Thiele et al. (12). In addition to the organ-specific maintenance biomass reactions, these WBMs also contain a whole-body biomass reaction that is constrained to carry a flux of 1 mmol/day/person in FBA simulations. Here, we simulated these male and female WBMs under the Average American and Gluten-free diets and determined the maximum flux of the sIEC maintenance biomass reaction in the WBMs with or without constraining the whole-body biomass reaction flux. We then calculated the relative percentage of the sIEC maintenance biomass reaction flux when constraining the whole-body biomass reaction flux to 1 mmol/day/person compared to that without, i.e., 100(sIEC maintenance biomass reaction flux in the WBM with the whole-body biomass reaction flux constrained at 1)/(sIEC maintenance biomass reaction flux in the WBM without constraining the whole-body biomass reaction flux). This resulted in 34.56% and 61.39% for the male WBM under the Average American and Gluten-free diets, respectively, and 99.88% or 78.67% for female under the same diets. We performed similar calculations when minimizing the sIEC maintenance biomass reaction within the WBMs, but the minimum values turned out to be the same as maximum values. These percentages were calculated to properly constrain the biomass reaction flux in sIEC GEMs for this study. To this end, we first calculated the maximum biomass reaction flux for sIEC GEMs constrained by transcriptional data under each diet. Next, the percentage obtained from the male or female WBM was multiplied by each IEC GEM’s maximum biomass flux under their respective diet, and the resulting value was set as the model’s biomass reaction flux. Calculating biomass flux separately for male and female samples thus allowed us to seamlessly incorporate sex as a biological variable in our analyses.

Computational simulation of GEMs was then performed using the COBRA Toolbox (9) in Python 3.8. For each metabolic task tested, its respective reaction flux was maximized while also minimizing deviations from gene expression data imposed using the soft constraints on reaction fluxes as noted above (McCreery and Zomorrodi, manuscript under preparation). The detailed optimization formulation is provided in the **Supplementary Text.**

### Shadow Price Analysis

Shadow prices in the context of linear optimization problems, such as those encountered in FBA for GEMs represent the impact of marginal changes in the right-hand side of the constraints on the objective function value. In the context of the sIEC GEMs for this study, the number of non-zero shadow prices represent the model’s capacity to adjust metabolic fluxes to optimize the activity of a metabolic task, which is used as the objective function. A greater number of non-zero shadow prices within a GEM indicates a higher flexibility of the network to adapt to perturbations. Shadow price analysis was performed using the COBRA Toolbox in Python 3.8. After performing FBA for each metabolic task, the total number of metabolites with non-zero shadow prices were recorded.

### Principal Component Analysis (PCA)

The optimal fluxes of the essential metabolic tasks were used as features for PCA (i.e., 59 features), while each of the 42 subjects served as samples (42 samples). To verify the status of the active CeD subject that is positioned far from the rest of the points in the PCA plot (shown in **Figure 1D**), we standardized the data along the first and second principal components (PC1 and PC2) independently using z-score transformation. The standardized values were then visualized to inspect the positioning of the active CeD subject relative to other subjects. PCA was performed using the *SKLearn* package in Python.

### Statistical analyses

Statistical hypothesis testing was conducted using the Mann-Whitney U (Wilcoxon Rank Sum) test using the *mannwhitneyu* function in Python’s *scipy.stats* package. All raw p-values were adjusted for multiple testing when relevant based on the Holm-Sidak method using the *multipletests* function in Python’s *statsmodels* package. Given the limitted sample size, significance was determined based on a somewhat lenient adjusted p-value threshold of 0.2.

### Drug target analysis

Genes corresponding to metabolic tasks with significant differential activity were extracted from the reactions’ gene-protein-reaction (GPR) rules in sIEC GEMs. These genes were then searched within the DrugBank database (66) to identify drugs that can up- or down-regulate their expression. Flux Coupling Analysis was performed using Flux Coupling Finder 2 (FCF2) v0.95b in MATLAB 2023b (3, 65).

## Acknowledgement

The authors would like to extend their gratitude to Dr. Maureen M. Leonard for sharing the gene expression count data related to reference (11) and Dr. Assieh Saadatpour for valuable insights during the data analysis phase.

## Author contributions

ARZ conceived the study and, together with CVM, interpreted the results and drafted the manuscript. CVM performed all analyses. DA conducted the whole-body model simulations. KM designed the list of food items for the average gluten-free diet. AF provided critical feedback on the results and manuscript. All authors have read and approved the final manuscript.

## Supplementary information

**Supplementary Table 1.** Diet formulation for the average American and gluten-free diets.

**Supplementary Table 2.** List of essential metabolic tasks for sIECs.

**Supplementary Table 3.** Standardized PCA coordinates (as shown in Supplementary Figure 1).

**Supplementary Table 4.** FBA results for the sIEC essential metabolic tasks.

**Supplementary Table 5.** Metabolite secretion fluxes into the bloodstream and intestinal lumen.

**Supplementary Table 6.** Non-zero shadow prices count for the essential metabolic tasks.

**Supplementary Table 7.** DrugBank Database search results for genes involved in the essential sIEC metabolic tasks.

**Supplementary Table 8.** Individual food items used for the construction of the average American diet.

**Supplementary Table 9.** Individual food items used for the construction of the gluten free diet.

**Supplementary Figure 1.** PCA with standardized data.

**Supplementary Figure 2.** Differential expression analysis of genes related to the essential metabolic tasks with differential activity.

**Supplementary Figure 3.** Essential metabolic tasks with non-significant differences in activity.

## Notes

### Competing Interest Statement

The authors have declared no competing interest.

