## Supplementary figures and images for "Investigating intestinal epithelium metabolic dysfunction in Celiac Disease using personalized genome-scale models"

### Supplemental Figure 1

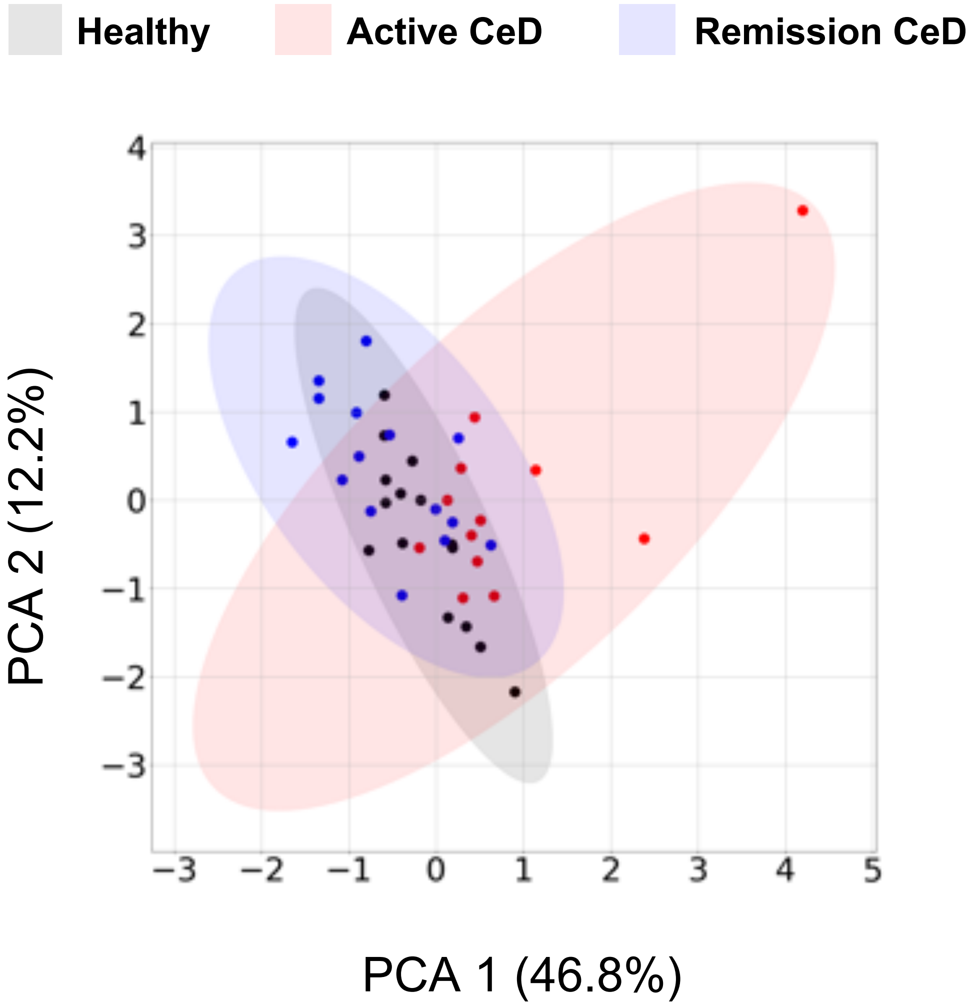


**Supplementary Figure 1.** PCA with standardized data.
