## Supplemental Figure 2 for "Investigating intestinal epithelium metabolic dysfunction in Celiac Disease using personalized genome-scale models"

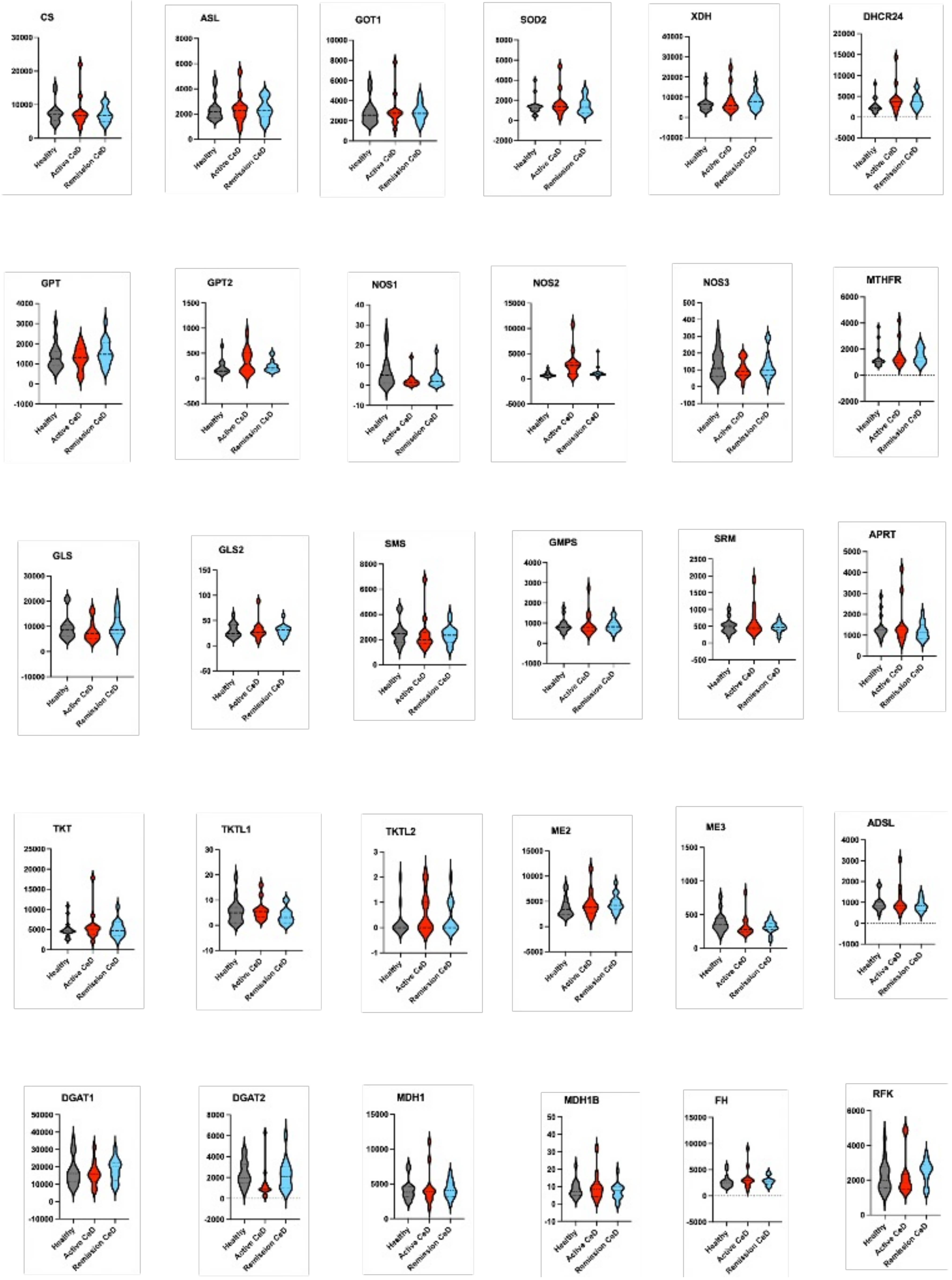


**Supplementary Figure 2.** Differential expression analysis of genes related to the essential metabolic tasks with differential activity.
