## Supplemental Figure 3 for "Investigating intestinal epithelium metabolic dysfunction in Celiac Disease using personalized genome-scale models"

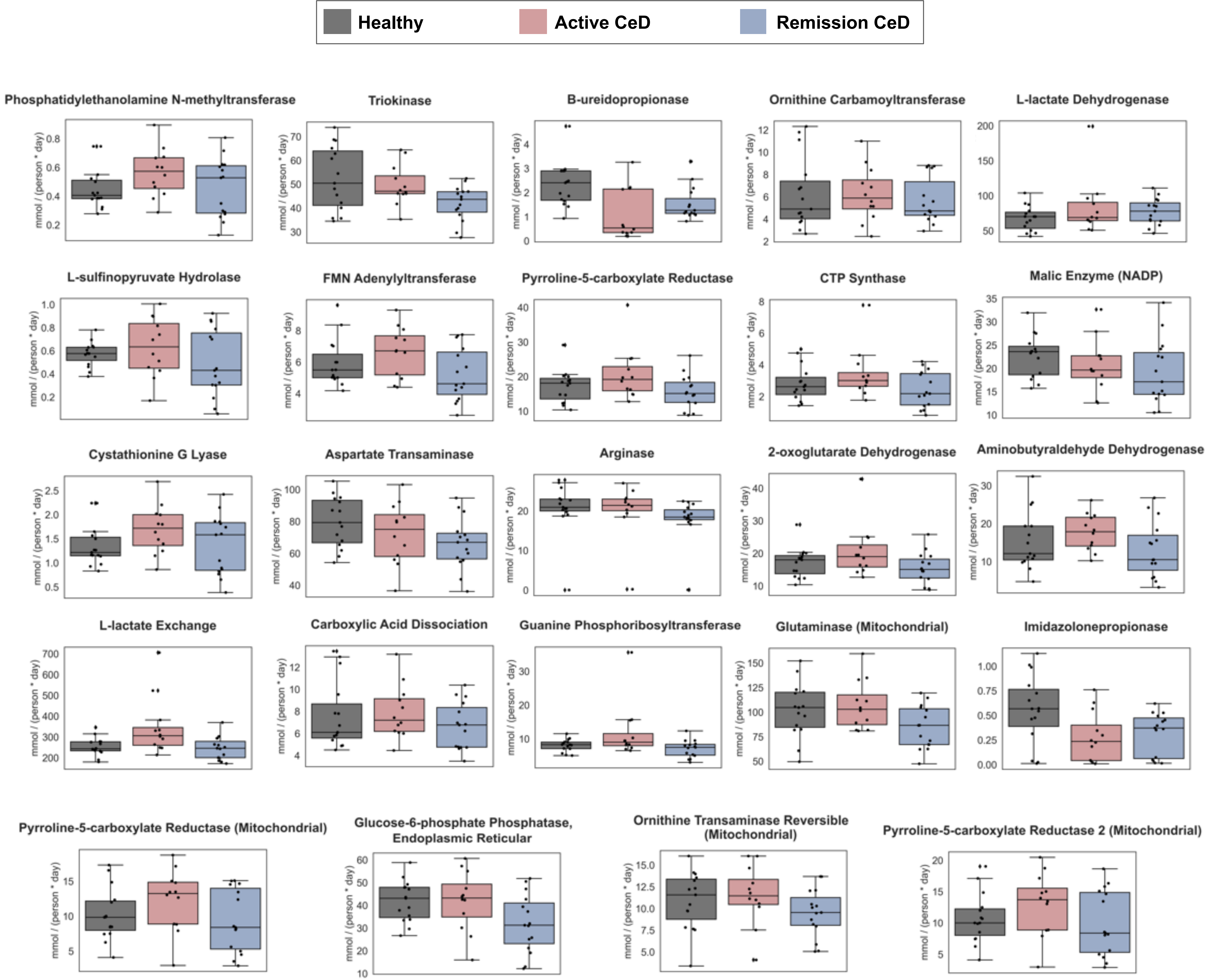


**Supplementary Figure 3.** Essential metabolic tasks with non-significant differences in activity.
